# Increased Inflammation as well as Decreased Endoplasmic Reticulum Stress and Translation Differentiate Pancreatic Islets of Pre-symptomatic Stage 1 Type 1 Diabetes and Non-diabetic Cases

**DOI:** 10.1101/2024.09.13.612933

**Authors:** Adam C. Swensen, Paul D. Piehowski, Jing Chen, X’avia Y. Chan, Shane S. Kelly, Vladislav A. Petyuk, Ronald J. Moore, Lith Nasif, Elizabeth A. Butterworth, Mark A. Atkinson, Rohit N. Kulkarni, Martha Campbell-Thompson, Clayton E. Mathews, Wei-Jun Qian

**Author notes:** Correspondence to: Wei-Jun Qian Clayton E. Mathews. These authors contributed equally to this work (Adam C. Swensen, Paul D. Piehowski, Jing Chen).

## Abstract

**Aims/hypothesis:** Progression to type 1 diabetes (T1D) is associated with genetic factors, the presence of autoantibodies, and a decline in β cell insulin secretion in response to glucose. Very little is known regarding the molecular changes that occur in human insulin-secreting β-cells prior to the onset of T1D. Herein, we applied an unbiased proteomics approach to identify changes in proteins and potential mechanisms of islet dysfunction in islet autoantibody-positive organ donors with pre-symptomatic stage 1 T1D (HbA1c ≤ 6). We aimed to identify pathways in islets that are indicative of β-cell dysfunction.

**Methods:** Multiple islet sections were collected through laser microdissection of frozen pancreatic tissues of organ donors positive for islet autoantibodies (AAb+, n=5), compared to age/sex-matched nondiabetic controls (ND, n=5) obtained from the Network for Pancreatic Organ donors with Diabetes (nPOD). Islet sections were subjected to mass spectrometry-based proteomics and analyzed with label-free quantification followed by pathway and functional annotations.

**Results:** Analyses resulted in ∼4,500 proteins identified with low false discovery rate (FDR) <1%, with 2,165 proteins reliably quantified in every islet sample. We observed large inter-donor variations that presented a challenge for statistical analysis of proteome changes between donor groups. We therefore focused on the three multiple AAb+ cases (mAAb+) with high genetic risk and their three matched controls for a final statistical analysis. Approximately 10% of the proteins (n=202) were significantly different between mAAb+ cases versus ND. The significant alterations clustered around major functions for upregulation in the immune response and glycolysis, and downregulation in endoplasmic reticulum (ER) stress response as well as protein translation and synthesis. The observed proteome changes were further supported by several independent published datasets, including proteomics dataset from *in vitro* proinflammatory cytokine-treated human islets and single cell RNA-seq data sets from AAb+ cases.

**Conclusion/interpretation:** In-situ human islet proteome alterations at the stage 1 of AAb+ T1D centered around several major functional categories, including an expected increase in immune response genes (elevated antigen presentation / HLA), with decreases in protein synthesis and ER stress response, as well as compensatory metabolic response. The dataset serves as a proteomics resource for future studies on β cell changes during T1D progression and pathogenesis.

## Introduction

Prior to the onset of type 1 diabetes (T1D), an asymptomatic period exists where ongoing autoimmunity and pancreatic islet β-cell dysfunction are present [1–3]. A staging concept of T1D pathogenesis has been proposed [2] with stage 1 being defined as individuals with genetic susceptibility, β-cell autoimmunity based on the presence of two or more islet autoantibodies (AAb+), along with normoglycemia [4, 5]. In Stage 2, AAb+ individuals progress to dysglycemia followed by Stage 3 or clinical onset of T1D. Studies over the past 6 decades have observed a decline of first-phase insulin release (FPIR) in response specifically to glucose in those subjects that progress to Stage 3 [3, 6–10]. Further, this defect in β-cell function has been observed years before clinical onset of T1D progressors in longitudinal studies with subjects starting at stage 0 or 1 [11, 12] as well as in those at risk for T1D development using cross sectional study designs [3, 13]. While genetics, autoimmunity, and β-cell dysfunction are all markers of T1D risk, gaps exist in our knowledge of the fundamental mechanisms leading to immune dysregulation as well as the basis for β-cell functional decline. Our understanding of pathogenic processes in human T1D were first elaborated using autopsy samples [14, 15] and have recently seen significant advances from the analyses of pancreatic tissues undertaken in collaborative networks, including Network of Pancreatic Organ donors with Diabetes (nPOD) [16]. Histological analysis has put forth the notion that selective β-cell loss occurs through islet infiltration of immune cells such as CD8^+^ and CD4^+^ T cells and CD20^+^ B cells within the islets (i.e., insulitis) [17, 18]. However, studies assessing β-cell loss and insulitis in pre-diabetic autoantibody-positive organ donors with risk-HLA have not observed significant declines β-cell mass and have noted the rarity of insulitis [18–20]. Interestingly, islets in non-diabetic AAb+ organ donors with risk-HLA haplotypes have been observed to have hyper-expression of HLA class I [21] and alterations in insulin prohormone processing [22]. To address the knowledge gaps, the current study used an unbiased proteomics approach to define molecular features of islets in the Stage 1 phase of T1D (mAAb+, genetic risk, and HbA1c<6) to gain a better understanding of the changes in β-cells during pre-T1D and identify potential molecular mechanism(s) associated with the initiation and progression of β-cell dysfunction in early stages of T1D.

The application of advanced ‘omics methodologies for broad molecular characterization of human islets represents a promising approach for obtaining detailed mechanistic information on the β-cell functional decline during Stage 1 T1D. Genomic and transcriptomic [23, 24], epigenetic [25], and proteomic [26] approaches have been applied to whole pancreas, isolated islets, and in some cases to single cells following islet cell isolation and sorting [27–29]. Mass spectrometry (MS)-based proteomics represents a promising global profiling technique for studying proteins or posttranslational modifications (PTMs). Herein, we aimed to identify in situ proteome alterations of the human islets at the stage 1 T1D through MS-based proteomics profiling of human islets by laser microdissection (LMD) of islet sections from AAb+ and age/sex matched control cases followed by LC-MS/MS analyses.

## Methods

### Laser microdissection (LMD) of human Islet Sections

Organ donors were selected with T1D-relevant autoantibodies (AAb+: against Insulin, GAD65, ZnT8, and IA-2A) from nPOD and pancreatic tissue sections were obtained (**Table 1**). Five AAb+ cases were selected (2 single AAb+ and 3 mAAb+). Five non-diabetic autoantibody-negative (ND) cases were selected matched in age and sex for each AAb+ donor creating 5 matched pairs (**Table 1**). Ten µm sections from fresh frozen OCT blocks were placed on PEN slides for LMD using previously described methods [30]. Sections were dehydrated in 100% methanol for 1 min, rinsed with DEPC-treated water, followed by incubation for 1 min in 75% ethanol, 1 min in 95% ethanol, and two 1 min washes in 100% ethanol. All solutions were prepared fresh on the day of use and stored on ice. Following the last 100% ethanol incubation, slides were blotted, placed in a RNAase-free desiccator for 8 mins at room temperature under vacuum, and immediately subjected to LMD using a Leica LMD7000 microscope. Islets were identified by inherent autofluorescence and morphology within exocrine regions. Dissections were performed using a 10X objective using the following settings: speed 5, balance 15, head correct 100, less 4755, and offset 50. Approximately 100 islet sections were obtained from each 10 µm tissue slide, corresponding to 1-2 islet equivalents (IEQs) and LMD sections were collected by gravity into the cap of a 0.5 mL LoBind microcentrifuge tube (Eppendorf, Enfield, CT, USA) for proteomic profiling. Two pools of ∼100 islet sections were obtained for each case. Samples were immediately frozen on dry ice then stored at -80°C until shipment on dry ice to PNNL.

**Table 1.**
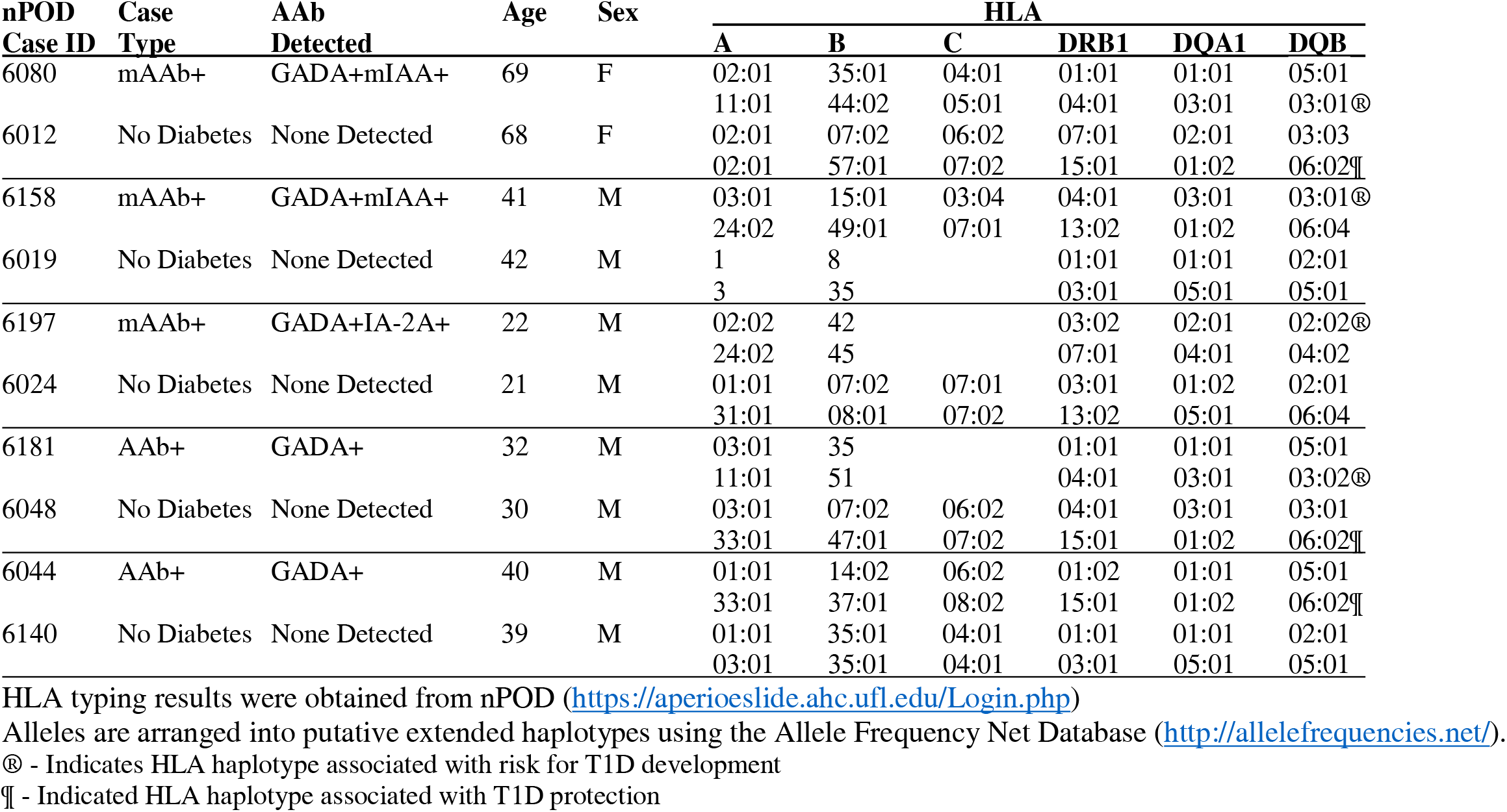
Patient Demographic Age and sex matched patient samples obtained from the Network for Pancreatic Organ Donors with Diabetes (nPOD).

| nPOD<br>Case ID | Case<br>Type | AAb<br>Detected | Age | Sex | HLA |  |  |  |  |  |
| --- | --- | --- | --- | --- | --- | --- | --- | --- | --- | --- |
|  |  |  |  |  | A | B | C | DRB1 | DQA1 | DQB |
| 6080 | mAAb+ | GADA+mIAA+ | 69 | F | 02:01<br>11:01 | 35:01<br>44:02 | 04:01<br>05:01 | 01:01<br>04:01 | 01:01<br>03:01 | 05:01<br>03:01® |
| 6012 | No Diabetes | None Detected | 68 | F | 02:01<br>02:01 | 07:02<br>57:01 | 06:02<br>07:02 | 07:01<br>15:01 | 02:01<br>01:02 | 03:03<br>06:02¶ |
| 6158 | mAAb+ | GADA+mIAA+ | 41 | M | 03:01<br>24:02 | 15:01<br>49:01 | 03:04<br>07:01 | 04:01<br>13:02 | 03:01<br>01:02 | 03:01®<br>06:04 |
| 6019 | No Diabetes | None Detected | 42 | M | 1<br>3 | 8<br>35 |  | 01:01<br>03:01 | 01:01<br>05:01 | 02:01<br>05:01 |
| 6197 | mAAb+ | GADA+IA-2A+ | 22 | M | 02:02<br>24:02 | 42<br>45 |  | 03:02<br>07:01 | 02:01<br>04:01 | 02:02®<br>04:02 |
| 6024 | No Diabetes | None Detected | 21 | M | 01:01<br>31:01 | 07:02<br>08:01 | 07:01<br>07:02 | 03:01<br>13:02 | 01:02<br>05:01 | 02:01<br>06:04 |
| 6181 | AAb+ | GADA+ | 32 | M | 03:01<br>11:01 | 35<br>51 |  | 01:01<br>04:01 | 01:01<br>03:01 | 05:01<br>03:02® |
| 6048 | No Diabetes | None Detected | 30 | M | 03:01<br>33:01 | 07:02<br>47:01 | 06:02<br>07:02 | 04:01<br>15:01 | 03:01<br>01:02 | 03:01<br>06:02¶ |
| 6044 | AAb+ | GADA+ | 40 | M | 01:01<br>33:01 | 14:02<br>37:01 | 06:02<br>08:02 | 01:02<br>15:01 | 01:01<br>01:02 | 05:01<br>06:02¶ |
| 6140 | No Diabetes | None Detected | 39 | M | 01:01<br>03:01 | 35:01<br>35:01 | 04:01<br>04:01 | 01:01<br>03:01 | 01:01<br>05:01 | 02:01<br>05:01 |

HLA typing results were obtained from nPOD (https://aperioeslide.ahc.ufl.edu/Login.php)
Alleles are arranged into putative extended haplotypes using the Allele Frequency Net Database (http://allelefrequencys.net/).
® - Indicates HLA haplotype associated with risk for T1D development
¶ - Indicated HLA haplotype associated with T1D protection

### Proteomic analysis of human islet sections

Microcentrifuge tubes containing dissected islets were centrifuged for 5 min at 21,000 rcf to pellet islet sections. Protein was extracted by adding 20 µL of homogenization buffer (8M urea, 50 mM Tris, 10 mM DTT, pH 8) followed by 10 pulses at 30% power in a UTR200 ultrasonic processor (Hielsher, Teltow, Germany). The extract was then incubated at 60°C for 30 min with 750 rpm shaking to promote protein denaturation and reduction. Samples were then transferred to low retention LC-MS vials (Waters Corporation, Milford, MA, USA) using 2 aliquots of 20 µL 50 mM Tris pH 8, creating a total volume of 60 μL for analyses on a simplified nanoproteomics platform [31] with online immobilized trypsin digestion and solid phase extraction followed by liquid chromatography (LC)-MS as previously described .

The simplified nanoproteomics platform was equipped with a LEAP autosampler allowing multiple samples to be queued and stored at 4°C prior to injection. Protein digestion was accomplished by an online immobilized enzyme reactor (IMER), a 150 μm i.d. column packed with immobilized trypsin beads. Eluent was then captured on a C18 trap column followed by LC-MS/MS. A 5-hour LC gradient length was applied using an in-house packed 50 μm i.d. fused silica capillary columns 75 cm in length. The system was coupled to an Orbitrap Q-Exactive Plus mass spectrometer (ThermoFisher Scientific) using a custom ESI interface comprised of 3 cm, chemically etched emitters coupled to the LC column using stainless steel unions (VICI Valco, Houston, TX, USA). Mass spectra were collected from 400– 2,000 m/z at a resolution of 70 K followed by data dependent HCD MS/MS at a resolution of 17.5 K for the ten most abundant ions.

### Multiplex immunofluorescence imaging

Multiplex immunofluorescence imaging of islet hormones (INS, GCG, SST), and the selected marker (NPTX2) was performed using methods previously described [18]. Formaldehyde-fixed paraffin embedded (FFPE) pancreas sections were obtained from nPOD. Sections were deparaffinized and rinsed in phosphate buffered saline (PBS) prior to high pH antigen retrieval. Blocking was carried out for 1 hour at room temperature in 10% normal serum. The antibody staining workflow is shown in **Suppl. Figure 1A**, and all primary and secondary antibodies are listed in **Suppl. Table 1**. Rabbit anti-human NPTX2 antibody (Sigma, HPA049799) was validated by the vendor using western blot and immunohistochemistry. After antibody staining, nuclei were stained using Hoescht S769121 (Nuclear Yellow, Abcam ab138903). Images were acquired using a Zeiss 710 confocal microscope. Islet images were acquired using a Zeiss 710 confocal microscope and 20x objective (Plan-Apochromat, 0.8 NA, 0.12µm/pixel, 8-bit depth with 16 frame averaging).

### Data analysis

Raw data from the mass spectrometer were analyzed in MaxQuant [32], version 1.5.3.30. with a protein level false discovery rate set at 0.01 with match between runs (MBR) enabled. Ion mass tolerances was set at 20 and 4.5 ppm for the first and main peptide searches. The quantification was performed by label-free quantification (LFQ) using default parameters. Pathway and function analyses were completed using Qiagen Ingenuity Pathway Analysis (IPA) software [33] with default parameters. Gene Ontology (GO) analyses utilized DAVID Bioinformatics Database Gene Functional Classification tool [34]. Visualization of significantly enriched gene ontology relationships was completed using Reduce and Visualize Gene Ontology (REVIGO) [35] web server with small (0.5) restrictive list generation using separate up- and down-regulated gene lists.

## RESULTS

### Comparative Proteome Profiling of Islets from AAb+ and Control Cases

Five AAb+ cases (3 mAAb+ and 2 single AAb+) and their age/sex matched ND controls were acquired from nPOD (**Table 1**). Samples were analyzed with an analytical workflow coupling LMD with the simplified nanoproteomics platform (**Figure 1A**) [31]. Relatively high reproducibility of the overall workflow was observed based on two replicates of islet samples from each donor (i.e., LMD islet sections from two adjacent tissue slides of the same donor) (**Figure 1B**).

**Figure 1.**
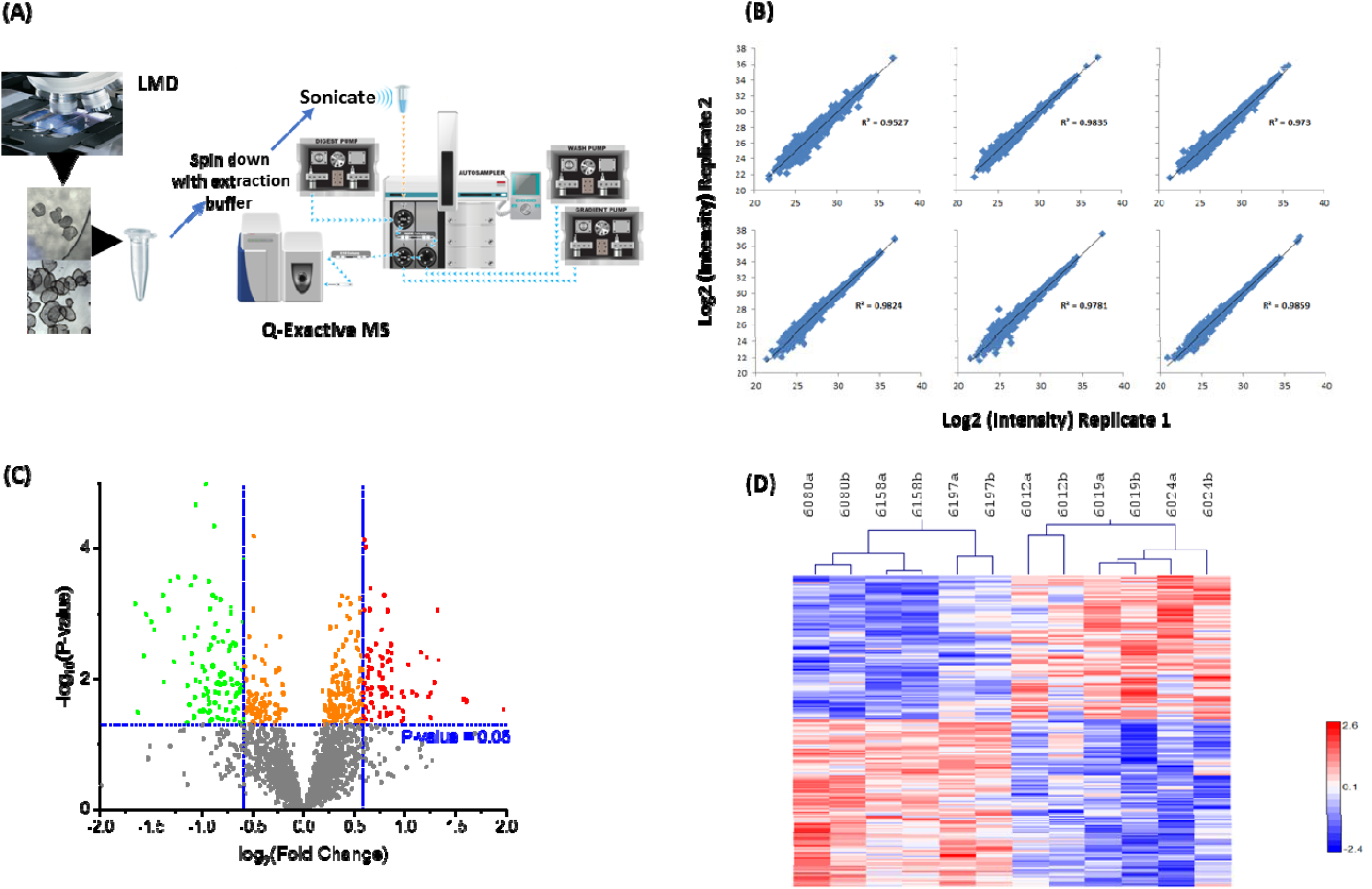
Experimental workflow, reproducibility, and overall significance (A) Experimental workflow. Two pools of ∼100 islet sections are collected per nPOD case via LMD from10 µm tissue sections. Proteins from each pool are extracted and subjected to online trypsin digestion and LC-MS/MS analysis. (B) Two pools of islet sections from each case were assessed as biological replicates for reproducibility. (C) Volcano plot of significant proteins between controls and stage 1 T1D patients. (D) Hierarchal heat map clustering of significant proteins for double AAb+ patients vs. controls (“a” and “b” denote biological replicates).

In total, ∼4,500 islet proteins were identified with <1% overall FDR (**Suppl. Table 2**), reflecting a significant coverage of the human islet proteome. The MaxQuant Match Between Runs (MBR) algorithm [32] was applied for label-free quantification analysis and 2,165 proteins were quantifiable with abundance values across biological samples (**Suppl. Table 3)**. Next, we tested for islet proteome differences between AAb+ (n=5) vs. ND samples (n=5). Principal component analysis (PCA) of the overall protein abundance data did not show significant differences between AAb+ and non-diabetic cases (data not shown). One reason for this lack of clear separation may be related to the limitation of classification of pre-T1D cases based solely on autoantibody positivity. Since the risk of T1D development is significantly greater in those with ≥2 AAb compared (>80% develop T1D) when compared to cases with a single autoantibody (<20% develop T1D) [4], we modified the analysis criteria to compare the mAAb+ cases with their age/sex matched ND controls. With this comparison (n=3 per group), 202 proteins were differentially expressed between mAAb+ and ND using a mixed effect model test where the FDR was controlled to be less than 0.15 and p value <0.025 (**Figure 1C and 1D, Suppl. Table 4**). The heatmap of **Figure 1D** shows clustering of mAAb+ versus ND cases based on the 202 proteins with differential expression. These data provide molecular evidence that *in situ* islet proteome change in Stage 1 individuals with mAAb^+^ [2].

### Altered Pathways and Functions Potentially Involved in Islet Dysfunction

To identify pathways and functions relevant to islet dysfunction in stage 1 T1D, the differential protein expression data were subjected to Ingenuity Pathway Analysis (IPA) [33]. Significant canonical pathways and cellular functions were altered in the islet proteome (**Figures 2 and 3, Suppl. Tables 5and 6**). These significant pathways and function categories center on several major classes including: 1) reduced translation or protein synthesis (EIF2 signaling, translation, tRNA charging) (**Figure 3A**); 2) increased catabolism (glycolysis, gluconeogenesis) **(Figure 3B)**; 3) reduced ER stress response (UPR) (**Figure 3C**); 4) increased immune response (antigen presentation pathway, viral infection) (**Figure 3D**); 4) increased cell death. Identification of several of these pathways or functions was confirmed by an independent informatics analysis based on Gene Ontology data (**Suppl. Figure 2**) using DAVID Gene Ontology Enrichment and visualized using REVIGO. For example, reduced translation (e.g., lower protein localization to ER, protein folding, and translation initiation), increased catabolism, and increased immune system response were also observed as significant categories.

**Figure 2.**
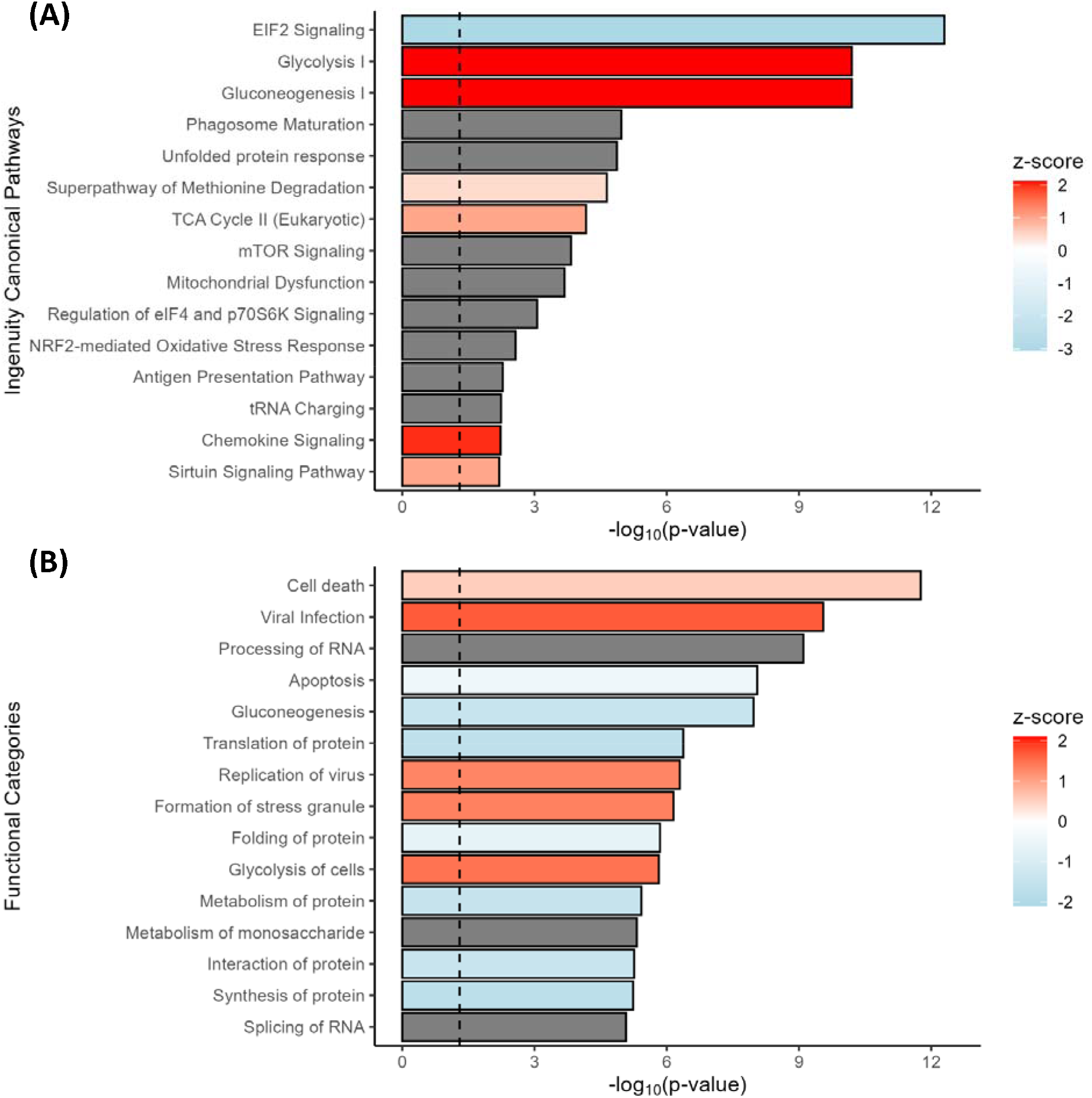
Significant enriched pathways and functional categories based on IPA analysis. Significant canonical pathways. Top 15 significant pathways were plotted with -log(p-value) along with Z-scores indicating predicted up-regulation (red) and down-regulation (blue) B) Functional categories. Top 15 significant functions were shown along with Z-score. The gray bars indicate no predicted Z-score values.

**Figure 3.**
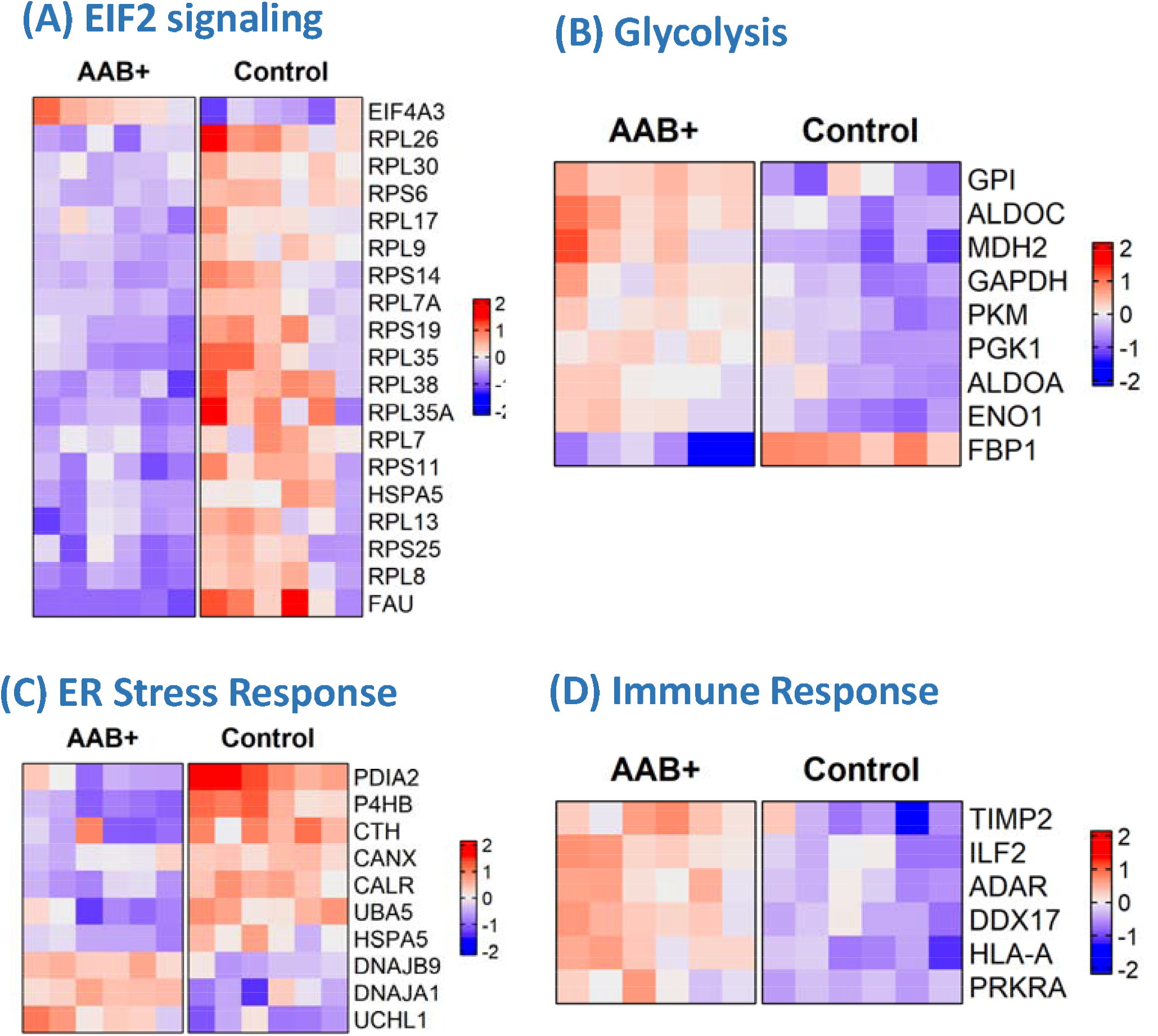
Heatmap of protein abundances in selected pathways. Heatmaps are displayed in relative protein abundances in a zero-centered log2 scale.

#### Deficiency in Translation and Protein Synthesis

Translation and protein synthesis were among the most significant altered biological processes in islets of the mAAb+ cases. A cluster of ribosomal proteins involved in protein synthesis, including 40S ribosomal proteins RPS6, RPS11, RPS14, RPS19, RPS25, FAU (a RPS30-ubiquitin fusion protein), as well as the 60s ribosomal proteins RPL7, RPL7A, RPL8, RPL9, RPL17, RPL26, RPL30, RPL35, RPL35A, and RPL38 were all observed to be reduced (**Figure 3A)**. This defect in protein synthesis occurs in islets early during T1D pathogenesis. A decrease in translation as well as short protein half-lives were reported as common features of protein regulation under viral infection, interferon exposure, or oxidative stress [36, 37], providing a potential biological link for these observations.

#### Increased Glycolysis

Interestingly, glycolysis and gluconeogenesis were observed as the second most significant pathways (**Figures 2 and 3B**). Most of the glycolytic enzymes (e.g., GPI: glucose-6-phosphate isomerase; ALDOA, ALDOC, GAPDH, PGK1, PKM, ENO) were observed as upregulated in mAAb+ islets; however, FBP1 (Fructose-1,6-bisphosphatase 1) was observed as significantly downregulated in mAAb+ islets.

#### Reduced ER Stress Response

The observation of reduced expression in the islet proteome included a decrease in several proteins related to ER stress response and UPR pathway (**Figure 3C).** PDI (P4HB), PDIA2, CALR and CANX are associated with ER function and involved in proper protein folding and quality control within the ER [38, 39]. DNAJA1, DNAJB9, and HSPA5 are heat shock protein family members directly involved binding to unfolded proteins, and HSP5A (or BiP) is known as a central regulator for ER stress. ER provides a quality-control system for the correct folding of proteins and for sensing stress, and the observed alterations suggest a potential failure of ER stress response within islets of mAAb+ cases.

#### Elevated Immune Response

Given the seroconversion and multiple autoantibody-positivity, we expected to observe an increased expression of immune response related proteins in mAAb+ islets. Proteins with significantly increased expression in mAAb+ islets (**Figure 3D**) included several variants of MHC class I cell surface antigens: HLA-A, PRKAR2B (a transcriptional regulator gene found in T-cells), ILF2 (a nuclear factor found in activated T-cells), and TIMP2 (a tissue specific matrix metalloprotease inhibitor involved in remodeling of the extracellular matrix). Hyperexpression of islet cell HLA class I antigens is reported as a defining feature of stage 1 T1D [21].

## DISCUSSION

Despite decades of research with clinical studies demonstrating a decline in β-cell function prior to diagnosis of autoimmune Type 1 diabetes, a clear molecular understanding of islet dysfunction and the pathways promoting loss of pancreatic β-cells in early stages of human T1D remains limited. While pathological evidence of insulitis and hyper-expression of HLA class I in AAb+ islets have been reported [18], the broad molecular changes at the proteome level of human islets at any pre-diabetic stage remains largely unknown. This lack of mechanistic understanding of islet dysfunction in humans is partly due to a lack of primary human tissue as well as the limited number of cases with pre-diabetes (mAAb+ with risk HLA). Here we utilize a highly sensitive technology to characterize the broad in situ changes in an unbiased molecular assessment with limited number of cases that have been collected.

The coupling of LMD and advanced-omics technologies is a timely direction that allows a detailed understanding of molecular changes in islets at Stage 1. The ability to perform broad measurements of protein abundances is a unique advantage of MS-based proteomics. While there are several recent reports on the application of proteomics to LMD isolated human islets [26, 40], the current study represents the first human islet proteome measurement from AAb+ cases, specifically mAAb+ cases with risk HLA. Although the present study can be considered as the exploratory stage due to the small number of Stage 1 cases (mAAb+ with risk HLA) that are available from the nPOD program, the overall work demonstrates clear potential value in performing detailed quantitative characterization of in situ islet proteome. The results represent an unbiased, realistic portrait of islet proteome changes during T1D progression. The availability of larger numbers of mAAb+ and recent-onset T1D cases (with residual β-cells) will allow an expansion towards a detailed mechanistic understanding of islet dysfunction.

It is important to note that the observed *in situ* islet proteome changes (based on both common pathways, functions, and proteins) were also consistent with several sets of orthogonal datasets, including isolated human islets treated *in vitro* with the combination of interferon gamma and interkeukin-1β [41] (**Figure 4A)**, and scRNA-seq dataset from isolated human islets from AAb^+^ cases and controls [27] (**Figure 4B)**. Globally, our data highlight biological processes that may play key roles in regulating islet dysfunction during the pre-diabetic stage, including increases in immune/inflammatory processes.

**Figure 4.**
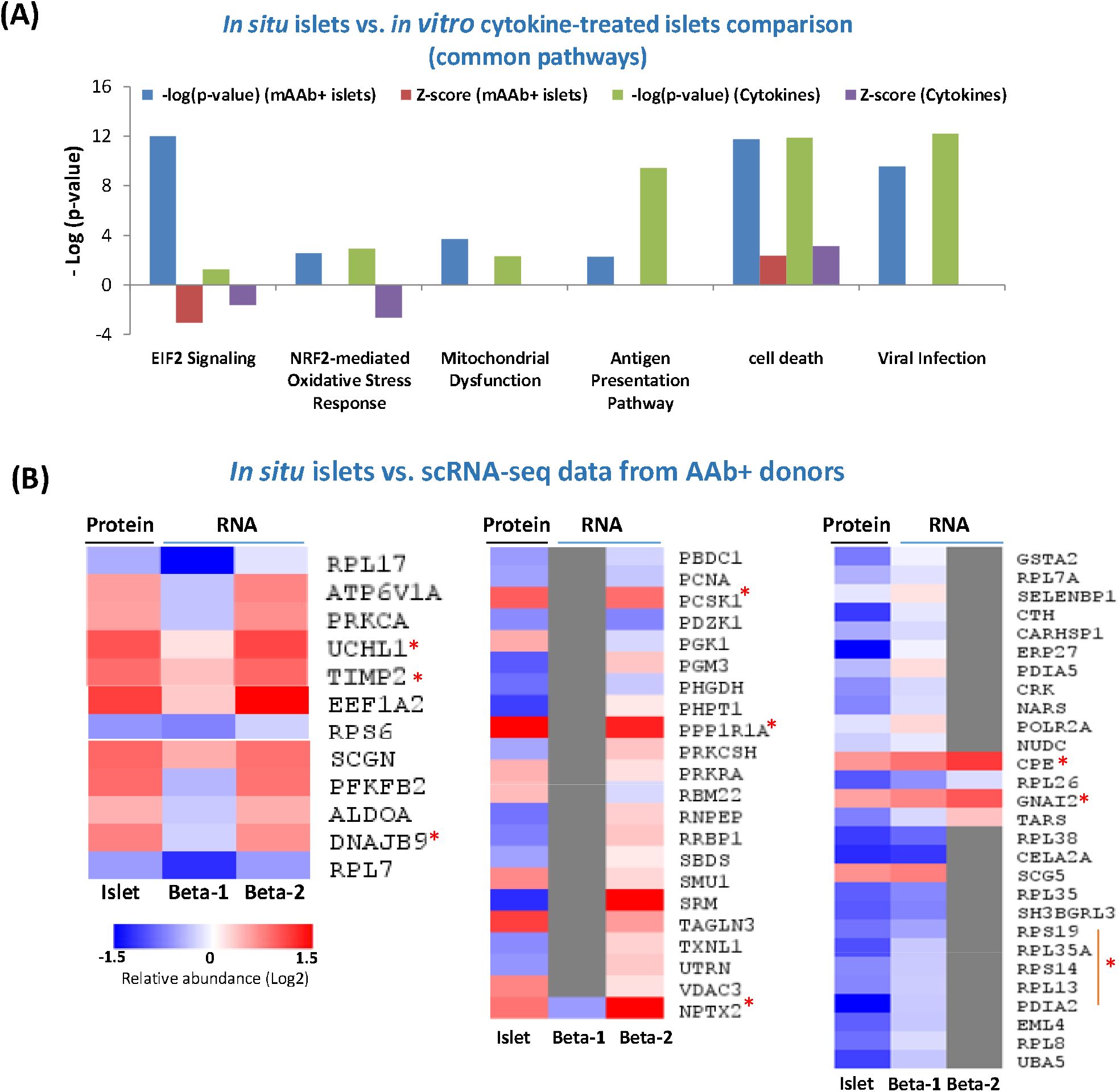
Comparisons of islet proteome changes with other independent datasets. (A) Pathway level enrichment comparison of our islet proteomics data with those from *in vitro* islets treated with cytokines (IFN-γ and IL-1β) (B) Comparison between islet proteome data with recent single cell RNA-seq data from β- cell subpopulations from the recent Human Pancreas Analysis Program (HPAP) study [27]. The list of significant proteins in the current study was mapped to those showing significant gene expression differences between AAb+ and control cases in the HPAP datasets. Selected protein/gene clusters are shown here. Red asterisks indicate selected proteins that were displayed in Figure 3 and a few additional proteins.

It is essential to note important differences as well, suggesting that *in vitro* models do not mirror the processes occurring in the islets during T1D pathogenesis. While a previous study reported an increase in CHOP (DDIT3) in both islet cells and islet-infiltrating immune cells it failed to demonstrate increases in PERK or IRE1 suggesting islets during T1D progress were not undergoing UPR or ER stress [42]. In the islets of Stage 1 T1D cases, our data point to a reduction in markers of the UPR with several major proteins in the ER quality control system being significantly down-regulated (**Figure 3B, Supplemental Figure 2**), including P4HB, PDIA2, CANX, CALR, and BiP. While our data did not detect the low-abundance UPR markers such as CHOP, PERK or IRE1, the observation of a significant downregulation of BiP (*HSPA5*), a UPR target gene product[43], provides evidence that UPR or ER stress response is not elevated in β-cells of at risk individuals [44] and, due to reductions in ER quality control proteins and BiP, may be inhibited [45]. The observed of a reduction of cystathionine γ-lyase (CTH) also provides an indication of reduced stress response [46]. While increased ER stress has been a postulated to be a component of β-cell failure in T1D, the data here and a previous publication suggest otherwise [42]. Interestingly, HLA class I upregulation in islets is a shared phenotype across individuals with T1D as well as Aab+ cases. Stimulation of UPR impairs the HLA class I-peptide complex presentation on the surface of cells [47–49]. Therefore, it is not surprising that the UPR pathway is reduced in mAAb+ risk-HLA islets with elevated HLA Class I (**Figure 3**).

The observed upregulation of hormone processing enzymes (PCSK1 and CPE) and proteins mediating insulin secretion (PPP1R1A) in the AAb+ pre-T1D stage is particularly interesting since altered β-cell prohormone processing and secretion has been increasingly recognized as an early and persistent finding in T1D [50]. PPP1R1A has been identified as down regulated in both T1D and T2D[51], and likely reflects a significant degree of β-cell loss due to the cell type specific expression of this protein. Another notable observation is the upregulation of neuronal pentraxin 2 (NPTX2), a secretory factor specifically expressed in β-cells within human pancreas as confirmed by multiplexed immunofluorescence (**Figure 5** **and Suppl. Figure 1**). The observed upregulation of NPTX2 is also consistent with recent scRNAseq data from human organ donors where NPTX2 upregulation was observed in the beta-2 subpopulation within the AAb+ cases (**Figure 4B**). NPTX2 has been reported to play a role in the regulation of anxiety and cognitive function [52, 53], but its role in islet biology is warranted for further study. Additionally, our observations include an upregulation in glycolytic enzymes (Figure 3).

**Figure 5.**
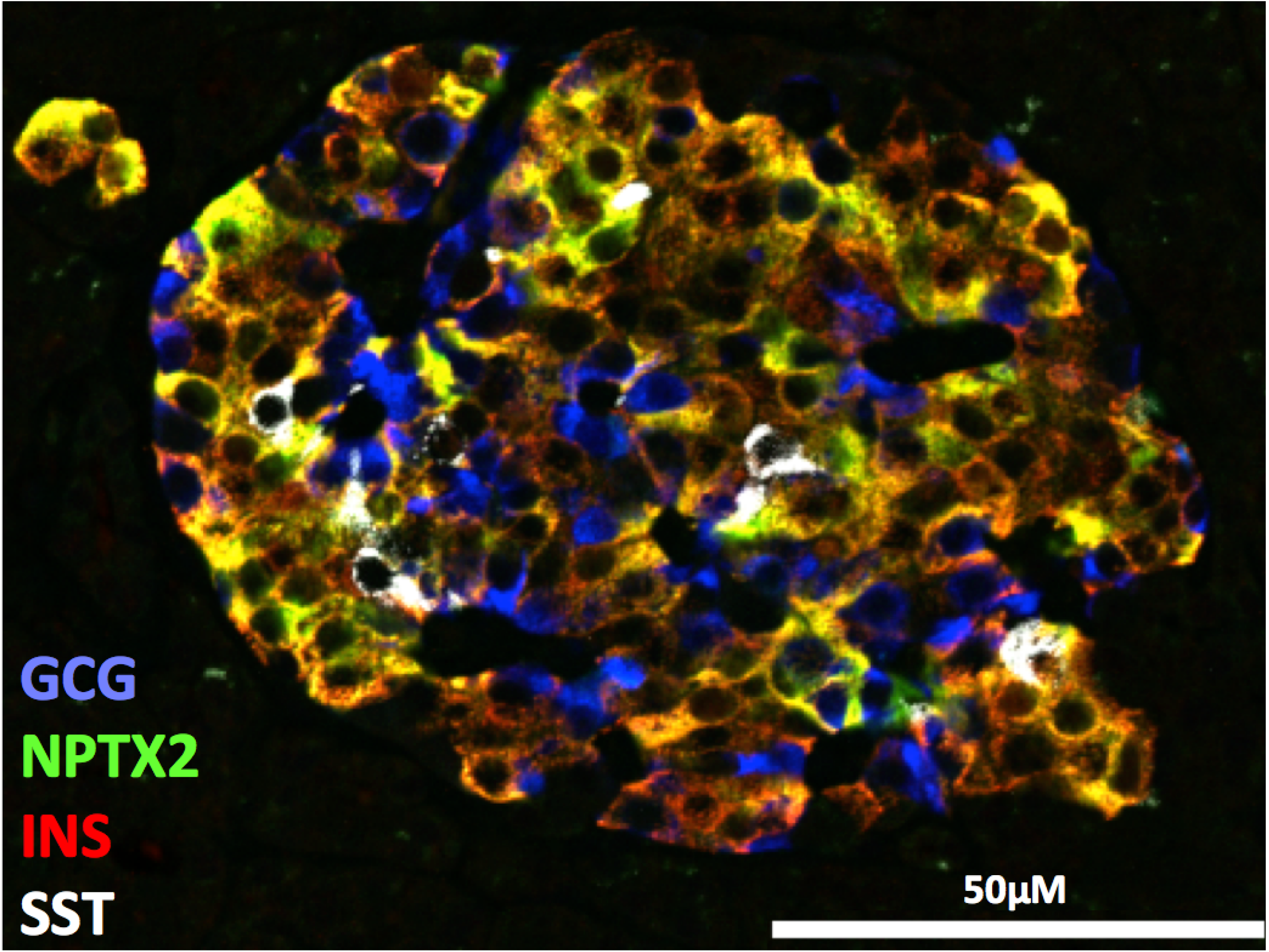
Immunolocalization of NPTX2 in human islets showed specific expression within β- cells. Multiplex immunofluorescence image of a representative islet is shown from a non- diabetic donor stained for GCG (blue), NPTX2 (green), INS (red, co-localization in orange), and SST (white).

These data provide evidence of metabolic abnormalities in stage 1 T1D [54] as well as dysregulation in insulin secretion since FBP1 was reported as an important regulator in insulin secretion [55]. Additional evidence of dysregulation of insulin secretion includes the observed upregulation of the G-protein coupled receptor associated protein GNAI2 in mAAb+ islets, a protein known to play a role in regulating insulin secretion [56]. Moreover, hyperactive glycolysis has been recently reported as a metabolic feature of β-cell dysfunction in aging and T2D [57].

In summary, quantitative profiling of the in-situ islet proteomes between mAAb+ and controls reveals that reduced translation and protein synthesis, failure of ER stress response, increased immune response and glycolysis are the major functional alterations in pre-diabetic islets. While our observations are corroborated by several orthogonal datasets, we acknowledge this study is limited by the number of available mAAb+ cases and future studies with larger numbers of cases will be warranted. The proteome datasets along with the protein candidates identified in the work could serve as a resource for generating new hypotheses for functional studies to identify the role of specific proteins in islet function that could also serve as potential targets for intervention.

## Supplementary Information

The online version contains peer-reviewed but unedited supplementary material available in the online version.

## Acknowledgements

We thank the donors and families of the donors for their invaluable contribution to our research and their help to further understand and hopefully cure T1D. This research was performed with the support of the Network for Pancreatic Organ Donors with Diabetes (nPOD), a collaborative T1D research project sponsored by JDRF. Organ Procurement Organizations (OPO) partnering with nPOD to provide research resources are listed at http://www.jdrfnpod.org/for-partners/npod-partners/. The proteomics work was performed in the Environmental Molecular Sciences Laboratory, a national scientific user facility sponsored by the DOE and located at Pacific Northwest National Laboratory, which is operated by Battelle Memorial Institute for the DOE under Contract DE-AC05-76RL0 1830. This work utilized a LEICA 7000 laser microdissection microscope purchased with an NIH shared instrumentation grant S10OD016350 and operated by the University of Florida Molecular Pathology Core (RRID:SCR_016601).

## Data availability

The LC-MS RAW datasets that support the findings of this study have been deposited in the online repository: MassIVE (https://massive.ucsd.edu/ProteoSAFe/static/massive.jsp) with accession MSV000090212.

## Author contributions

W.J.Q., C.E.M., R.N.K. and M.C.T. designed the experiments. J.C., P.D.P., A.C.S., X.Y.C, S.S.K, V.A.P., L.N., E.A.B. and R.J.M. performed different parts of experiments or analyzed the data. A.C.S., W.J.Q., C.E.M., M.C.T., and R.N.K. wrote the manuscripts. M.A.A. contributed to the study discussion and edited the manuscript.

## Research in context

### What is already known about this subject?

- The best-known biomarker for determining T1D risk continues to rely on the type and number of islet autoantibodies in people who are first degree relatives of a T1D patient. These autoantibodies generally appear years prior to the onset of dysglycemia and clinical presentation.
- First phase insulin-secretion is altered during stage 1 T1D despite the ability to maintain normoglycemia.
- Islet inflammation (insulitis) is present but uncommon in stage 1 T1D organ donors (observed in 2 out 18 donors or 11%).

## What is the key question?

- What alterations in the global proteome distinguish human islets from stage 1 T1D, which would provide mechanistic understanding of early stage of islet dysfunction?

## What are the new findings?

- 202 proteins were significantly altered in the in-situ islet proteome analyses between stage 1 T1D cases and matched controls.
- Dysregulated pathways in stage 1 T1D include reduced protein translation and ER stress response, and an increased islet immune response.

## How might this impact clinical practice in the foreseeable future?

- Our data serve as a resource for future functional studies aimed at developing interventions that target the dysregulated pathways in pancreatic β-cells.
- Proteins identified in our study may serve as biomarkers of stage 1 T1D.

## Abbreviations

LMD: Laser Micro-dissection
AAb: Autoantibody
ER: Endoplasmic Reticulum
UPR: Unfolded protein response
nPOD: Network for Pancreatic Organ Donors with Diabetes GADA Glutamic Acid Decarboxylase Autoantibodies
IAA: nsulin Autoantibodies
IA-2A: Insulinoma-2-associated Autoantibodies Zn-T8 Zinc Transporter 8 Autoantibodies

## Supplementary Information

**Supplemental Data 1.**: Supplemental Tables 1-6 in excel spreasheets including the list of total identified islet proteins, significantly altered proteins, and the list of all quantified proteins in human islets.

**Supplemental Figure 1.**
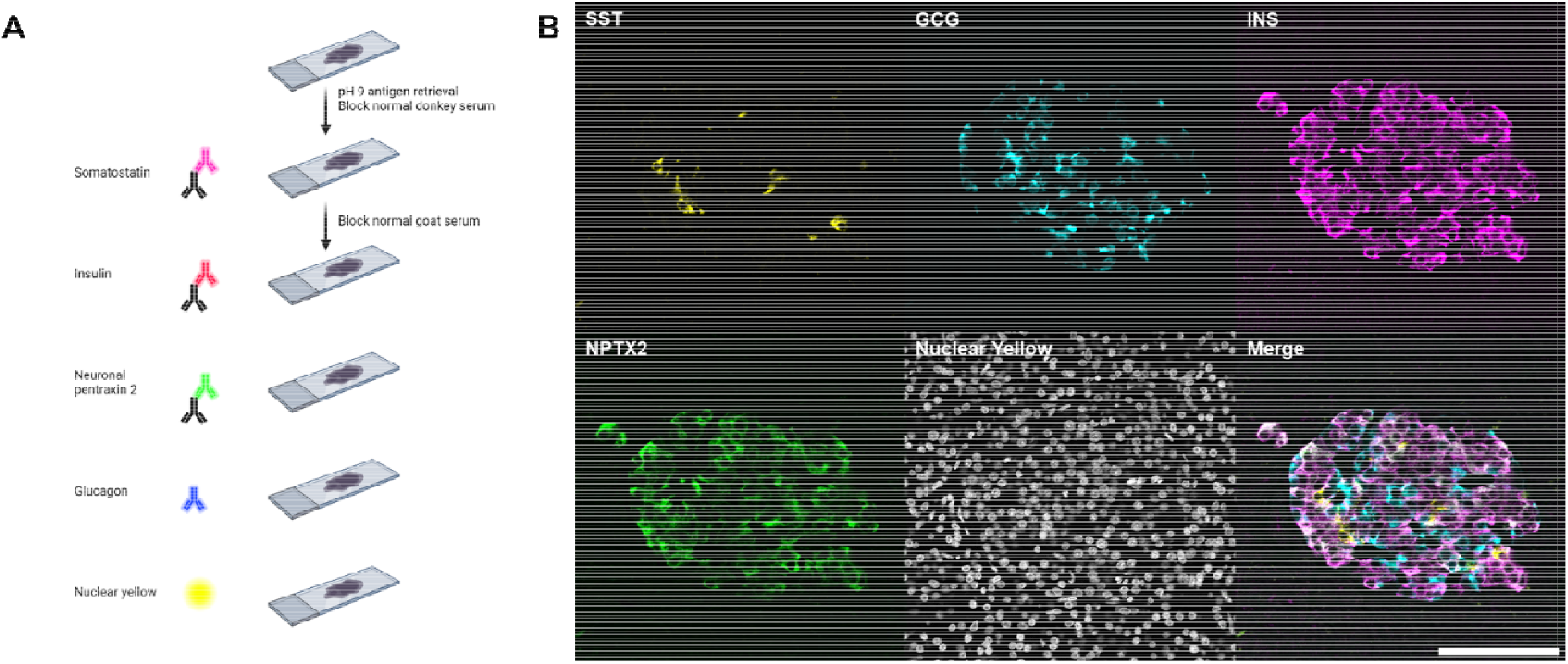
: Multiplex immunofluorescence imaging of islet proteins. (A) The antibody staining workflow; (B) Images of individual channel and merged image showing that NPTX2 is mainly expressed in beta cells within islets.

**Supplemental Figure 2.**
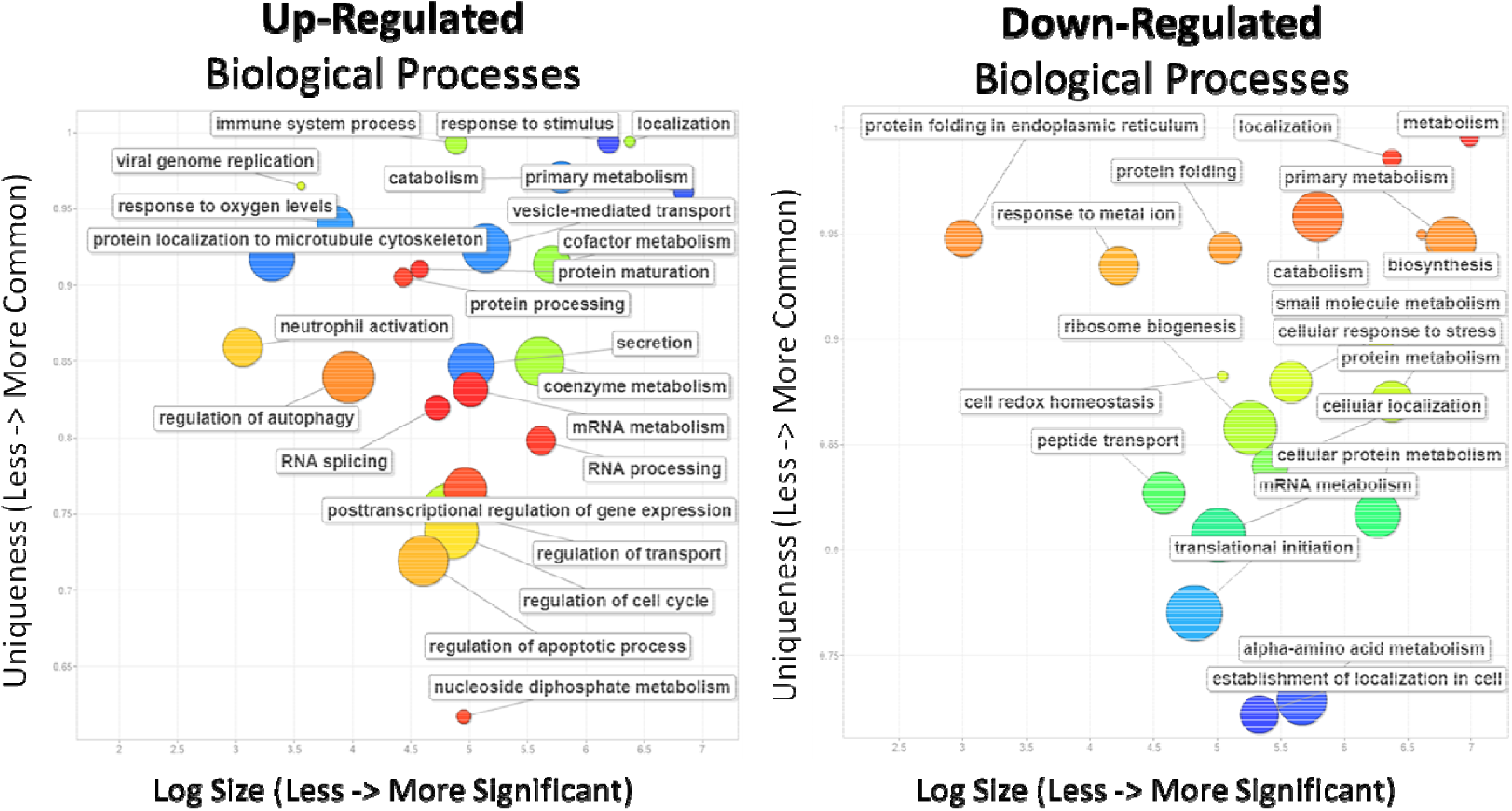
: Revigo visualization of ontology of up- and down-regulated biological processes. The size of the circle indicates the number of proteins identified in the process. The vertical position indicates the uniqueness of the pathway with the top being more common, while the signficance of th pathway (log size) increases to the right.

